# Cultivation of host-adapted *Cryptosporidium parvum* and *Cryptosporidium hominis* using enteroids for cryopreservation of isolates and transcriptomic studies of infection

**DOI:** 10.1101/2023.12.06.570384

**Authors:** Miner Deng, Tianyi Hou, Xinjie Mao, Jie Zhang, Fuxian Yang, Yanting Wei, Yongping Tang, Wanting Zeng, Wanyi Huang, Na Li, Yaoyu Feng, Lihua Xiao, Yaqiong Guo

**Affiliations:** State Key Laboratory for Animal Disease Control and Prevention, South China Agricultural University, Guangzhou, Guangdong, China

**Author notes:** Address correspondence to Lihua Xiao, or Yaqiong Guo,. Miner Deng and Tianyi Hou contributed equally to this work. The order was decided based on the fact that Miner Deng took a lead role in the performance of the project and preparation of the manuscript, while Tianyi Hou was responsible for the bioinformatic analysis and preparation the figures and tables.

**Keywords:** *Cryptosporidium*, organoids, culture, transcriptomics, host-pathogen interactions

## Abstract

*Cryptosporidium hominis* and *Cryptosporidium parvum* are major causes of severe diarrhea in humans. Comparative studies of them are hampered by the lack of effective cultivation and cryopreservation methods, especially for *C. hominis*. Here, we described adapted murine enteroids for the cultivation of one *C. parvum* IId subtype and nonhuman primate-adapted *C. hominis* Ib, Im, and In subtypes, which allowed the complete development of the pathogens, producing oocysts infectious to mice. Using the system, we developed a novel cryopreservation method for *Cryptosporidium* isolates. In comparative RNA-seq analyses of *C. hominis* cultures, the enteroid system generated significantly more transcriptomic responses of both pathogen and host genes than the conventional HCT-8 cell system. In particular, the infection was shown to upregulate PI3K-Akt, Wnt, Ras,TNF, NF-κB, IL-17, MAPK, and innate immunity signaling pathways and downregulate Wnt and Hippo signaling pathways, host cell metabolism, and parasites in enteroid cultures had significantly higher expression of genes involved in oocyst formation. Therefore, the new culture model provides a valuable tool for comparative studies of the biology of divergent *Cryptosporidium* species.

**IMPORTANCE**

The two dominant species for human cryptosporidiosis, *Cryptosporidium hominis* and *Cryptosporidium parvum*, differ significantly in host range and virulence. Up to date, biological studies of *Cryptosporidium* spp. are almost exclusively done with bovine-adapted IIa subtypes of *C. parvum*, which is the species with effective laboratory animal models and in vitro cultivation methods. Here, we describe modified procedures for the generation of murine enteroids for successful cultivation of both nonhuman primate-adapted *C. hominis* subtypes and a *C. parvum* IId subtype, producing oocysts infective to mice. In addition, we have developed a novel cryopreservation method using the system for long-term storage of *Cryptosporidium* isolates. RNA-seq analyses of *C. hominis* cultures indicate that the enteroid culture system generates host and pathogen transcriptomic responses similar to those in natural infection. This new development alleviates a technical bottleneck in cryptosporidiosis research, and provides an example for other difficult-to-culture pathogens of major public health importance.

---

Cryptosporidiosis is an important cause of severe diarrhea in humans, responsible for major waterborne epidemics in high-income countries and childhood malnutrition and death in low- and middle-income countries (1, 2). *Cryptosporidium hominis* and *Cryptosporidium parvum* are two main species responsible for human cryptosporidiosis (3). Between them, *C. hominis* has a narrow host range, being found in mainly equine animals in addition to humans and other primates, while *C. parvum* infects a broad range of animals including primates, ruminants, equine animals, and rodents (4). The two also differ significantly in genomics and virulence (5, 6).

Within the two major human-pathogenic species, there are significant variations in phenotypic traits at the subtype family level (4). Among the over 20 subtype families of *C. parvum*, IIa and IId are the two major zoonotic ones commonly found in humans and farm animals. Between them, IIa subtypes are common in intensively farmed dairy calves, causing outbreaks of zoonotic cryptosporidiosis in most high-income countries (4). In contrast, IId subtypes are more common in lambs and goat kids in Europe, Mideast countries and Oceania and in dairy calves in China and Sweden, causing mostly sporadic zoonotic infections in humans (7). Similarly, *C. hominis* subtype families differ in host preference and virulence. In addition to the common Ia, Ib, Id, Ie and If subtype families in humans, several new subtype families have been identified in equine animals (Ik) and nonhuman primates (Ii, Im, and In) (4).

Currently, biological studies on *Cryptosporidium* spp. have mostly concentrated on *C. parvum* IIa subtypes, while the IId subtypes prevalent in some countries are rarely used (8). In addition, studies of the biological characteristics of *C. hominis* are non-existent because of the lack of effective cultivation and easily accessible animal models (8).

Although several significant progresses, such as the development of potential treatments (9) and genetic modification tools (10), have been made in recent years, we still have a rudimentary understanding of *Cryptosporidium* spp. (11). One of the most important reasons is the lack of effective and economic culture platform to support the complete life cycle of *Cryptosporidium* spp. and propagation of oocysts in vitro.

Human adenocarcinoma cell lines (such as HCT-8 cells) are widely used in *in vitro* studies of *C. parvum*, but they support only partial development of the pathogen without the production of oocysts, the infective stage (12). To support complete the development of *C. parvum* and maintain long-term cultures, several studies have reported the establishment of various 3D culture systems. However, these novel culture systems are difficult to manipulate and require expensive and specialized equipment (12, 13). In addition, the HCT-8 cells used differ biologically from native epithelial cells in the intestine.

Organoids, products of the 3D culture of stem cells, retain the biological characteristics, unique markers and other functional properties of the native tissues (14). Recent studies indicate that organoids derived from intestinal crypts can be used in drug screening, organ transplant, modeling of genetic diseases, and studies of host-pathogen interactions (15). Heo *et al.* demonstrated that *C. parvum* could be cultured in human intestinal and lung organoids (16). However, the culture system developed requires the microinjection of individual organoids, hampering its routine applications. More recently, Wilke *et al.* developed a new enteroid system for the complete development of *C. parvum* (17). It grows stem cells digested from the ileum in Transwells and uses an air liquid interface (ALI) to induce cell differentiation. The new enteroid system sustains the growth of IIa subtypes of *C. parvum*, leading to the generation of infective oocysts. However, none of the new and conventional systems have been used effectively in the cultivation of host-adapted *C. parvum* IId subtype family and *C. hominis*.

Another major bottleneck of research progress is the lack of accessible cryopreservation of *Cryptosporidium* oocysts. Currently, researchers propagate *Cryptosporidium* spp. in laboratory animals every 2∼6 months, which is labor-intensive and expensive. Recently, a cryopreservation method of *C. parvum* by freezing permeabilized oocysts in silica microcapillaries was reported (18). To overcome the limited volume (∼2 μl oocyst suspension) of microcapillaries, the research group developed a modified cryopreservation protocol by using ‘vitrification cassettes’, and adapted it for the cryopreservation of *C. hominis* oocysts (19, 20). However, these methods require specialized equipment that is not available in most laboratories.

Here, we refined the procedures for the generation of murine enteroids and ALI cultures, including modifications of feeder cell induction, digestion of enteroids, and the medium used in the ALI culture. We have used the new ALI system in the cultivation of one *C. parvum* IId subtype and three nonhuman primate-adapted *C. hominis* subtypes. Based on the ALI culture system, we established a novel cryopreservation method of *C. parvum* for long-term maintenance of isolates. In addition, we compared transcriptomic responses in HCT-8 and ALI cultures infected with *C. hominis* through RNA-seq analysis. The new system supports the complete development of both *C. parvum* and *C. hominis*, permits the cryopreservation of

*Cryptosporidium* isolates, and allows comparative studies of the biology and host-pathogen interactions of multiple *Cryptosporidium* species and isolates in a controlled environment.

## RESULTS

### Development of an enteroid-derived ALI system for *C. parvum* cultivation

We generated enteroid stocks from intestinal crypts for establishing the ALI system (Fig. 1A). The intestinal crypts isolated from the intestines of C57BL/6 mice looked initially like pellets (Fig. 1B). After three days of culture, enteroids started to form, with budded crypt-like and villus-like domains (Fig. 1B). The enteroids generated could be maintained in culture for over 6 months, and cryopreserved for future usage. Through one or more rounds of passages, numerous enteroids with few single cells or mesenchymal cells could be produced for establishing the ALI culture system.

**FIG 1.**
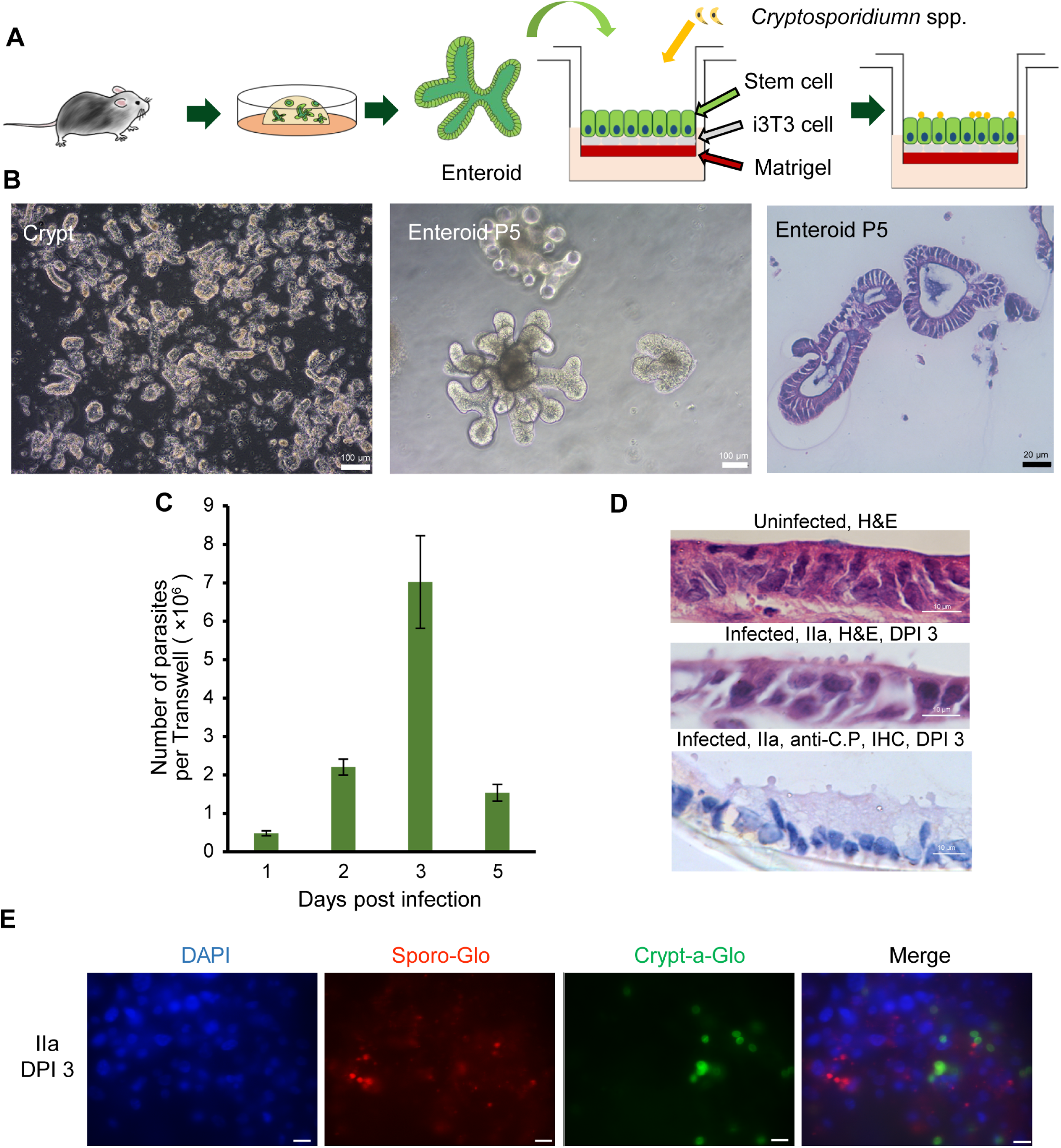
Establishment of an enteroid-derived air-liquid interface (ALI) system for cultivation of *Cryptosporidium parvum* IIa-Waterborne isolate in vitro. (A) Schematic outline of the process of creating mouse intestinal organoid stocks and establishing the ALI system for infection. (B) Images of murine intestinal crypts (left image), enteroid passage No. 5 (P5, middle image), and H&E-stained sections of P5 (right image). Crypts isolated from mouse intestines looked like pellets initially. After 3 days of cultivation, the enteroids formed with budding structures and an internal chamber. (C) Propagation of the *C. parvum* IIa subtype in the ALI system as measured by qPCR, n = 3. (D) Histological (H&E) and immunohistochemistry (IHC) microscopic images of the ALI system infected with *C. parvum* IIa-Waterborne oocysts at DPI 3. Developmental stages (globular structures) of *C. parvum* are visible on the surface of cells in infected ALI cultures. Scale bars, 10 μm. (E) Immunofluorescent microscopy of ALI monolayers infected with IIa-Waterborne sporozoites, showing the presence of developmental stages (stained with Sporo-Glo mAb) and oocysts (stained with Crypt-a-Glo mAb) at DPI 3. Scale bars, 10 μm.

Using the enteroids grown in Transwells, we adapted the ALI culture system following procedures described previously (17) with modifications of the medium used, the induction of feeding cells and the digestion of enteroids. The ALI cultures were infected with 2×10^5^ oocysts of the IIa-Waterborne isolate of *C. parvum* three days after the top medium was removed. Results of qPCR analysis showed that parasite growth was amplified by ∼70-fold from the initial infection dose. The parasite load peaked at DPI 3 (Fig. 1C). Histological examination of the ALI cultures revealed the presence of parasites on the surface of ALI monolayers (Fig. 1D). Immunofluorescent staining of the ALI monolayers infected with purified sporozoites using antibodies against developing stages (Sporo-Glo) and oocysts (Crypt-a-Glo) confirmed the production of oocysts in the ALI cultures (Fig. 1E).

### The ALI system allowed the full development of the IId subtype of *C. parvum* and production of infectious oocysts

To determine whether the ALI culture system would support the growth of host-adapted IId subtypes of *C. parvum*, we infected ALI cultures with the virulent IId-HLJ (2×10^5^ oocysts per Transwell). Compared to IIa-Waterborne, the growth of IId-HLJ was significantly higher with an amplification of ∼300-fold from the infection dose (Fig. 2A). Both H&E and immunohistochemical (IHC) analyses showed a large number of parasites on the surface of the ALI monolayers (Fig. 2B). Immunofluorescence microscopy using antibodies against cellular markers villin and Muc2 confirmed that the ALI treatment promoted cell differentiation (Fig. 2C). The production of oocysts was supported by immunofluorescence microscopy using antibodies against oocysts (Fig. 2D).

**FIG 2.**
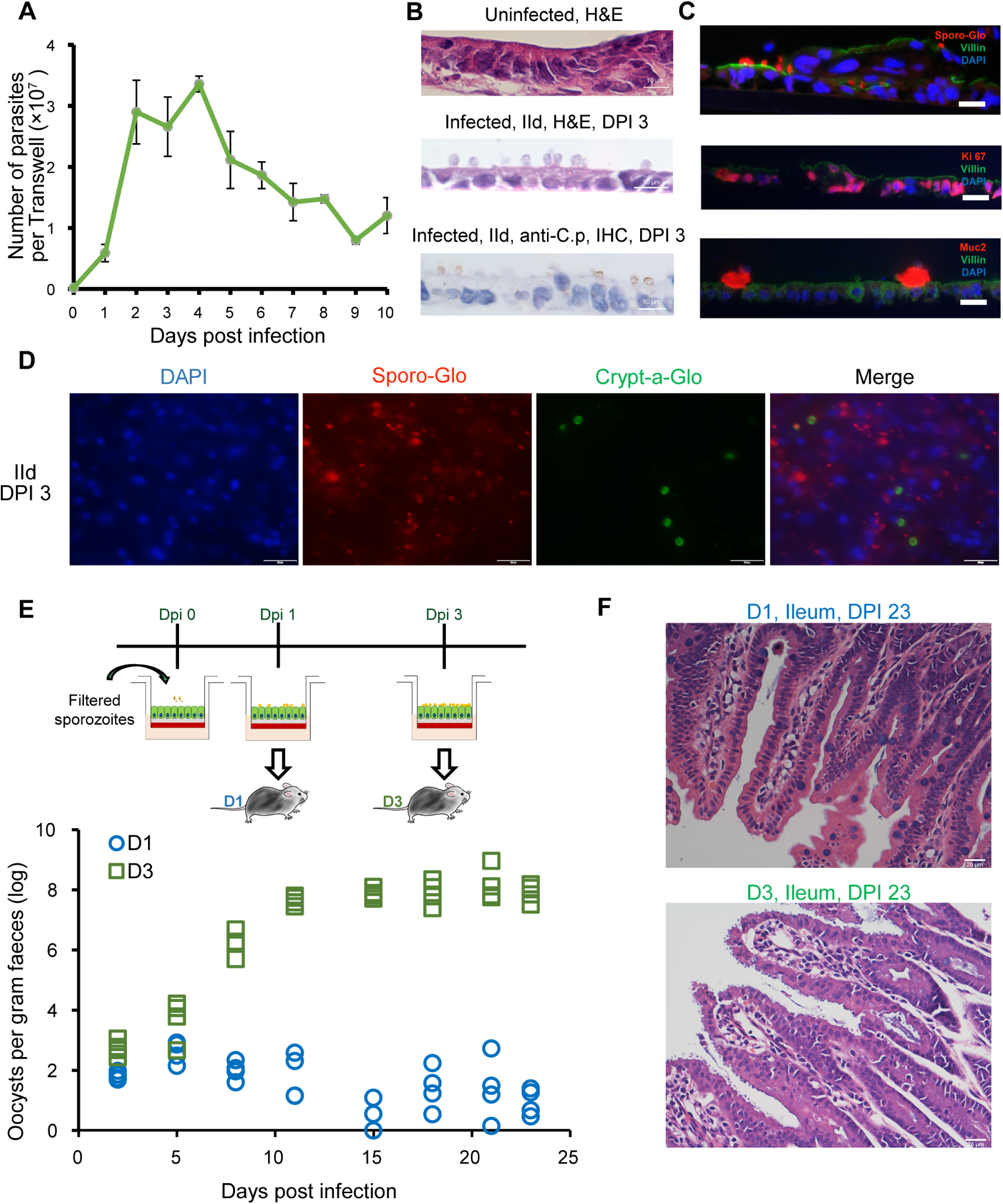
Complete development of the IId subtype of *Cryptosporidium parvum* in the murine organoid-derived air-liquid interface (ALI) system. (A) Proliferation of IId-HLJ isolate cultured in the ALI system as measured by qPCR, n = 3. (B) Images of ALI sections stained with H&E or rabbit antibodies against *C. parvum* using immunohistochemistry (IHC), DPI 3. Developmental stages of *C. parvum* are visible on the surface of cells in infected ALI cultures. Scale bars, 10 μm. (C) Immunofluorescence images of the sections of the ALI cultures infected with IId-HLJ at DPI 3 using antibodies against Ki-67 (a marker for replicating cells), Muc2 (a marker for goblet cells) and Sporo-Glo (reactive with intracellular stages of *C. parvum*). Scale bars, 10 μm. (D) Immunofluorescent microscopic images of the ALI cultures infected with IId-HLJ sporozoites after staining with Sporo-Glo (reactive with developing stages) and Crypt-a-Glo (reactive with oocysts) antibodies against *C. parvum*), DPI 3. Scale bars, 20 μm. (E) Patterns of oocyst shedding in mice (n = 4) orally gavaged with ALI cultures that were infected with IId-HLJ sporozoites for 1 (D1) and 3 (D3) days. The cultures were treated with 10% bleach to kill immature stages prior to mouse inoculation. (F) Images of H&E-stained sections of the small intestine from these mice at DPI 23. Developmental stages of *C. parvum* are visible only on the surface of the ileum from mice infected with D3 cultures. Scale bars, 20 μm.

To assess the infectivity of oocysts generated, we infected IFN-γ knockout (GKO) mice with parasites harvested from ALI cultures infected with sporozoites, after killing immature stages by bleach treatment of the harvested parasites (Fig. 2E). Mice infected with DPI 1 material (containing no oocysts) showed no clinical signs of cryptosporidiosis. In contrast, mice infected with DPI 3 material (containing newly generated oocysts) showed inactivity, rough hair, arched back and sticky fecal pellets, starting at DPI 7. The oocyst shedding reached peak at DPI 10, with oocyst per gram (OPG) of feces values reaching ∼10^8^ (Fig. 2E). Consistent with the oocyst shedding results, H&E-stained sections of intestinal tissues from mice infected with DPI 3 material showed the presence of high parasite burdens, while no parasites were seen in tissues from mice inoculated with DPI 1 material (Fig. 2F).

We performed scanning (SEM) and transmission electron microscopy (TEM) to further examine the development of *C. parvum* in ALI cultures. SEM images of ALI cultures infected with IId-HLJ showed the presence of parasites on the surface of the monolayers in clusters (Fig. 3A). Elongation and reduction in numbers of microvilli were seen in cells infected with the parasites (Fig. 3A). Multiple life cycle stages were observed under TEM, including meronts, macrogamonts and oocysts (Fig. 3B).

**FIG 3.**
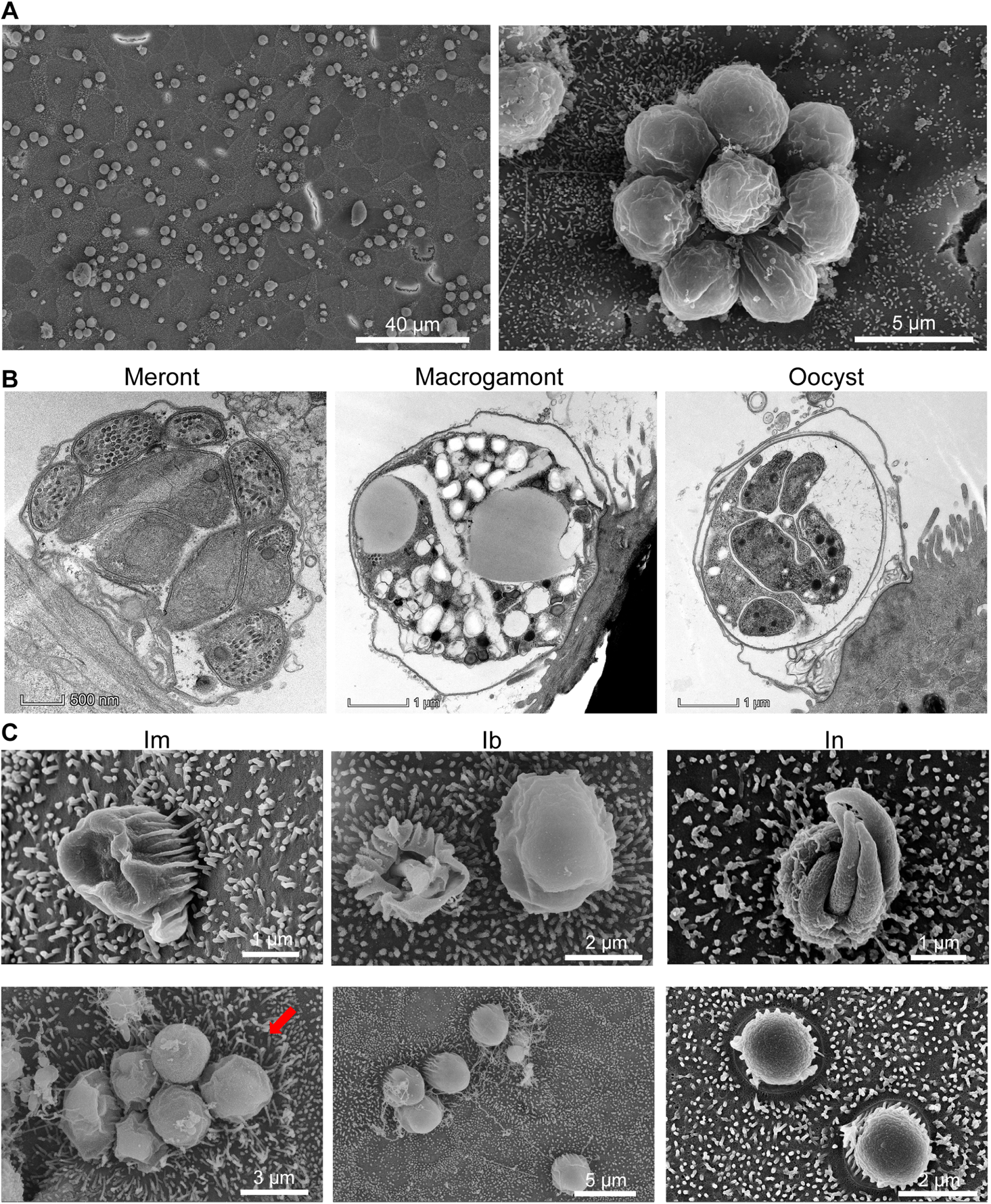
Scanning (SEM) and transmission electron microscopy (TEM) images of developmental stages of *Cryptosporidium* spp. in the enteroid-derived air-liquid interface (ALI) system. (A) SEM images showing the development of *C. parvum* IId-HLJ isolate on top of the ALI monolayers. The left image shows the elongation and reduction in numbers of microvilli on infected cells, while the right image shows a cluster of parasites developed from eight merozoites released from one mature meront. (B) TEM images of one meront, macrogamont and oocyst each of *C. parvum* within the parasitophorous vacuole on the apical surface of ALI monolayers. The ALI cultures were infected with 3×10^5^ sporozoites per Transwell. (C) SEM images of *C. hominis* developed in ALI cultures for three days. The ALI monolayers were infected with Ib, Im and In isolates. Elongation and reduction in numbers of microvilli are visible near the developing parasites in the lower left image (indicated by a red arrow).

Sporulated oocysts surrounded by parasitophorous vacuole membrane (PVM) were recognized on the surface of some infected cells at DPI 3, confirming that the ALI system supported the full development of the *C. parvum* IId subtype (Fig. 3B).

### The ALI system permitted accessible cryopreservation of *C. parvum*

To examine whether ALI cultures infected with *C. parvum* could be cryopreserved, we froze ALI cultures 6 h (ALI-6h) or 12 h (ALI-12h) after infection with sporozoites of IId-HLJ (Fig. 4A). After storage in liquid nitrogen for over one month, the preserved cultures were thawed and seeded onto ALI cultures (Fig. 4A). Results of the qPCR analysis showed that the numbers of recovered parasites from cryopreserved ALI-6h cultures increased between DPI 1 and 3 (*P* = 0.0421), while the increase in parasite burden was much less obvious in ALI cultures inoculated with cryopreserved ALI-12h cultures (*P* = 0.8546) (Fig. 4B). Immunofluorescent examinations showed that both developing stages (stained with Sporo-Glo) and oocysts (stained with Crypt-a-Glo) were presented at DPI 3 in ALI cultures infected with recovered parasites (Fig. 4C). H&E-stained sections showed that parasites were adhering in the top of ALI monolayers infected with recovered ALI-6h cultures, although we failed in detecting parasites on the surface of cultures infected with cryopreserved ALI-12h cultures, probably due to the lower infection intensity (Fig. S1A). This was confirmed by the results of the IHC analysis of cultures (Fig. S1A).

**FIG 4.**
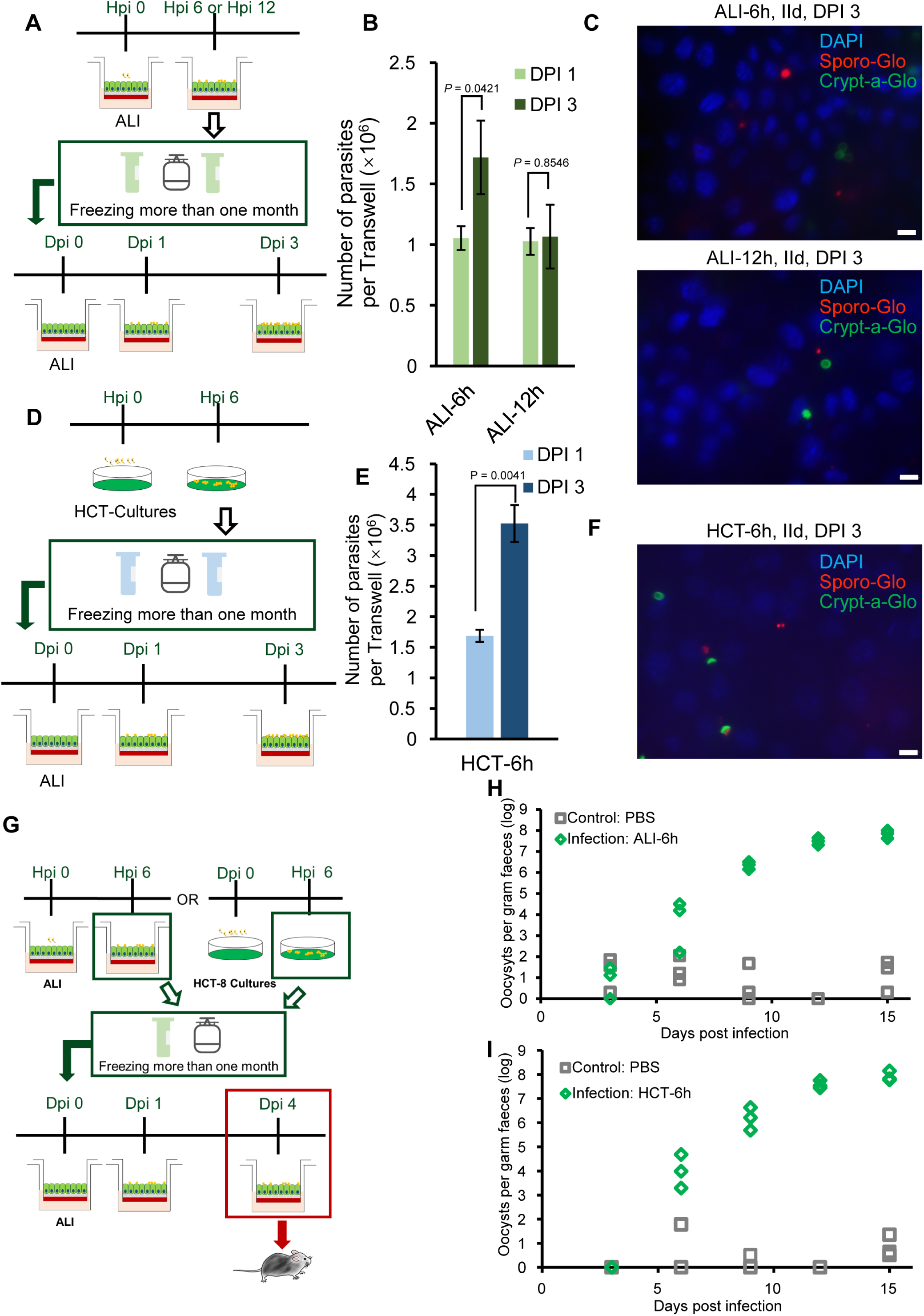
An accessible cryopreservation method of *Cryptosporidium parvum* using the air-liquid interface (ALI) system. (A) Schematic outline of the process of the cryopreservation of *C. parvum* using the ALI system. *C. parvum* IId-HLJ was cultured in ALI for 6 (ALI-6h) or 12 h (ALI-12h). The infected ALI cells were cryopreserved in liquid nitrogen for over one month prior to be inoculated into new ALI cultures for recovery and continued cultivation of *C. parvum* for one or three more days. (B) Proliferation of the recovered parasites in ALI cultures at DPI 1 and DPI 3, n = 3. New ALI cultures were infected with cryopreserved ALI-6h or ALI-12h. (C) Immunofluorescence images of the ALI monolayers infected with ALI-6h or ALI-12h for three days, showing the presence of developing stages (reactive to Sporo-Glo antibodies) and oocysts (reactive to Crypt-a-Glo antibodies). Scale bars, 10 μm. (D) Schematic outline of the process of the cryopreservation of *C. parvum* using inoculation of ALI cultures with parasites cultured in HCT-8 cells. New ALI cultures were infected with cryopreserved HCT-8 cells 6 h after they were infected with IId-HLJ (HCT-6h). (E) Proliferation of recovered parasites in ALI cultures one and three days after they were inoculated with cryopreserved HCT-6h, n = 3. (F) Immunofluorescence image of the ALI monolayers infected with HCT-6h for three days. Scale bars, 10 μm. (G) Schematic outline of evaluation of infectivity of oocysts produced in ALI cultures that were inoculated with cryopreserved ALI/HCT-8 cultures infected with *C. parvum* for 6 hours. (H) & (I) Patterns of *Cryptosporidium* oocyst shedding in mice. Infection: mice were orally gavaged with the ALI materials infected with ALI-6h (H) or HCT-6h (I) for four days; Control: mice were orally gavaged with PBS.

To assess if ALI infection with routine HCT-8 cultures of *C. parvum* would allow the cryopreservation of the pathogen, we performed ALI infection studies with cryopreserved HCT-8 cells that were infected with sporozoites of the IId subtype of *C. parvum* for 6 h (HCT-6h). The thawed HCT-6h cultures were seeded onto ALI cultures, with the parasite growth being determined by qPCR and immunofluorescent analysis (Fig. 4D). The parasite load in ALI cultured infected with cryopreserved *C. parvum* in HCT-6h increased ∼2-fold between DPI 1 and DPI 3 (*P* = 0.0041) (Fig. 4E). Both developing stages and oocysts were seen at DPI 3 in ALI cultures (Fig. 4F). Developing stages were also observed on the surface of ALI cultures infected with HCT-6h under H&E and IHC microscopy (Fig. S1B).

To determine if the oocysts collected from ALI cultures infected with cryopreserved *C. parvum* were viable and infectious, we inoculated GKO mice with the ALI cultures described above. Cryopreserved ALI-6h or HCT-6h were cultured further in ALI system for 4 days to produce oocysts. Mice inoculated with bleach-treated ALI materials started to shed oocysts at ∼ DPI 6, with peak oocyst shedding of ∼10^8^ OPG at DPI 12 (Fig. 4H and 4I). Subtyping of the parasites by sequence analysis of the *gp60* gene confirmed that the oocysts inoculated and produced after infection were identical. Microscopy of H&E-stained sections of the ileum from mice infected with HCT-6h or ALI-6h material showed heavy *C. parvum* infection (Fig. S1C).

In the present study, we used only frozen cultures that had been stored for one month. Future studies should evaluate the impact of longer storage on the revival of cryopreserved parasites.

### The ALI system supported complete development of multiple *C. hominis* isolates

To assess the infectivity of *C. hominis* isolates to murine enteroids, we infected ALI cultures with 1×10^5^ oocysts (Fig. 5A and 5B) or 4×10^5^ sporozoites (Fig. 5C & 5D) per Transwell. The three genetically unique *C. hominis* subtypes from macaque monkeys were used, including Ib, In and Im. They showed a similar infection pattern in ALI cultures, with the parasite burden peaking at DPI 3 (Fig. 5A). Among them, Ib and In generated peak parasite burden of over 1.5×10^7^ parasites per Transwell, while Im generated peak parasite burden of 4×10^6^ (Fig. 5A). For all three *C. hominis* isolates, the number of parasites per Transwell at DPI 3 was significantly higher than that at DPI 1 (Ib, *P* = 0.0047; In, *P* = 0.0130; Im, *P* = 0.0513) (Fig. 5A).

**FIG 5.**
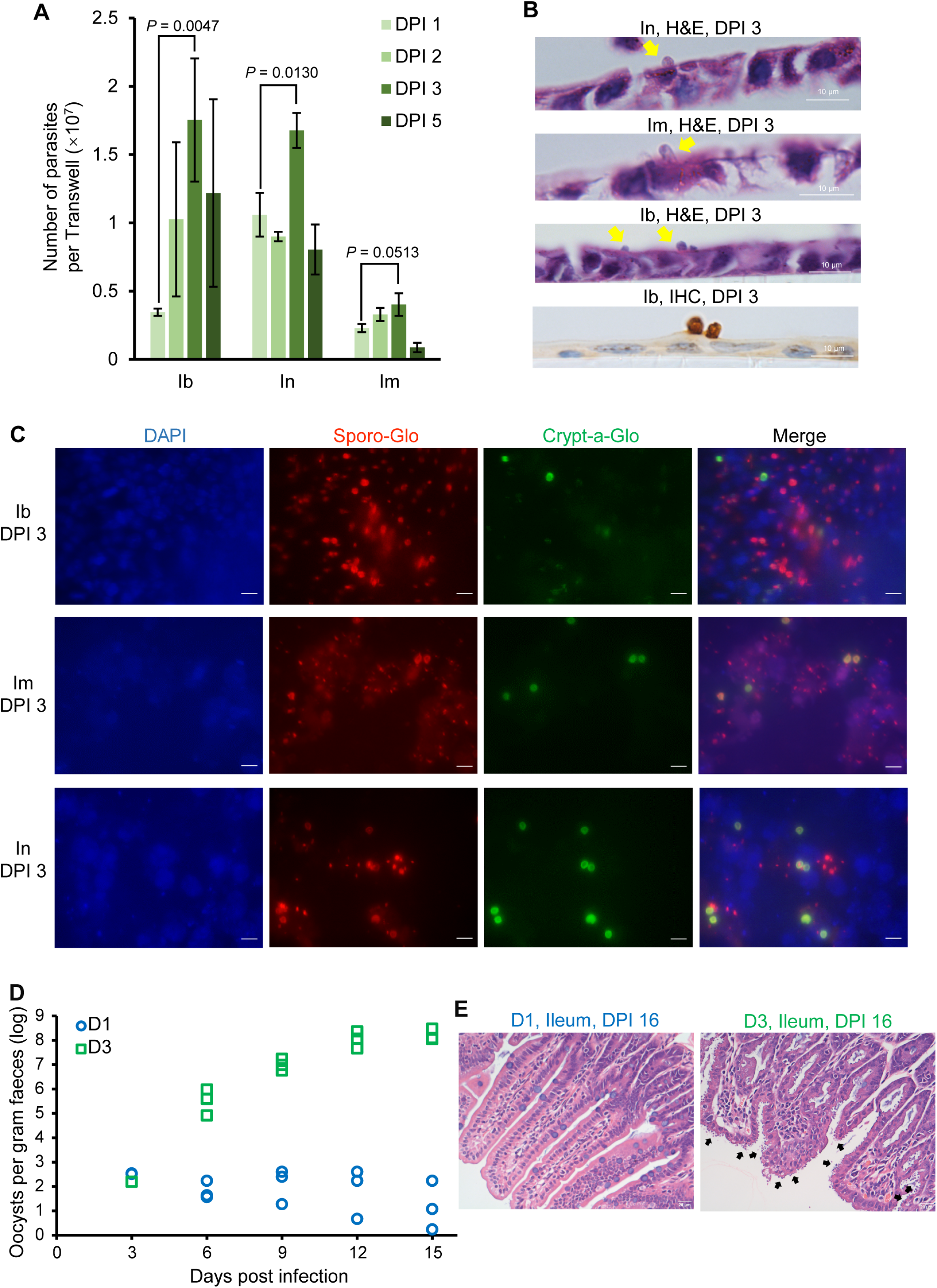
Cultivation of *Cryptosporidium hominis* using the enteroid-based air-liquid interface (ALI) system. (A) Growth of Ib, Im, and Im subtypes of *C. hominis* in ALI after infection with 2×10^5^ oocysts/Transwell, n = 3. (B) H&E microscopy images of ALI monolayers infected with Ib, Im and In, showing the presence of parasites (marked with arrows) on the surface of epithelial cells. (C) Immunofluorescence images of ALI cultures infected with 4×10^5^ sporozoites of three *C. hominis* subtypes, showing the presence of developmental stages (reactive to Sporo-Glo antibodies) and oocysts (reactive to Crypt-a-Glo antibodies). (D) Patterns of oocyst shedding in mice (n = 4) orally gavaged with ALI materials that were infected with sporozoites of *C. hominis* Ib subtype for one (D1) and three days (D3). Scale bars, 10 μm. (E) Images of H&E-stained sections of the ileum of mice infected with D1 or D3 materials described above for 16 days. Scale bars, 20 μm.

H&E microscopy revealed the presence of numerous parasites on the surface of the ALI monolayers (Fig. 5B). Immunofluorescence microscopy of the ALI cultures infected with sporozoites showed the presence of both developing stages (reactive to Sporo-Glo antibodies) and mature oocysts (reactive to Crypt-a-Glo antibodies) (Fig. 5C). SEM analysis of the ALI monolayers identified the presence of parasites on the surface of many cells, with apparent elongation of microvilli near the parasites (Fig. 3C).

To determine whether the *C. hominis* oocysts produced in ALI cultures were viable and infectious, we infected mice with ALI cultures that were infected with filtered 5×10^5^ Ib sporozoites per Transwell and harvested at DPI 1 or DPI 3. Mice infected with DPI 3 materials started shedding oocysts at DPI 6 and reached peak at DPI 15 with OPG values of ∼10^8^, while mice infected with DPI 1 material remained uninfected (Fig. 5D). The Ib subtype identity of parasites shed by mice was confirmed by *gp60* sequence analysis. Microscopy of H&E-stained sections of the ileum from mice infected with DPI 3 cultures showed numerous parasites on the surface of intestinal epithelial cells, while those inoculated with DPI 1 culture remained uninfected (Fig. 5E).

### Transcriptomic responses induced by *C. hominis* differed significantly between ALI and HCT-8 cultures

To characterize host cell responses to *C. hominis* infection in vitro, we analyzed the transcriptomes of ALI and HCT-8 cultures infected with the Ib subtype of *C. hominis* at DPI 3. Principle component analysis (PCA) showed significant differences in host gene expression between the infected and control groups in both ALI and HCT-8 cultures (Fig. 6A). A total of 4141 differentially expressed genes (DEGs) were identified in murine enteroids infected with *C. hominis* at DPI 3, with 2328 up- and 1813 down-regulated genes, while 2141 DEGs were identified in HCT-8 cultures with 701 up- and 1440 down-regulated genes (Fig. 6B).

**FIG 6.**
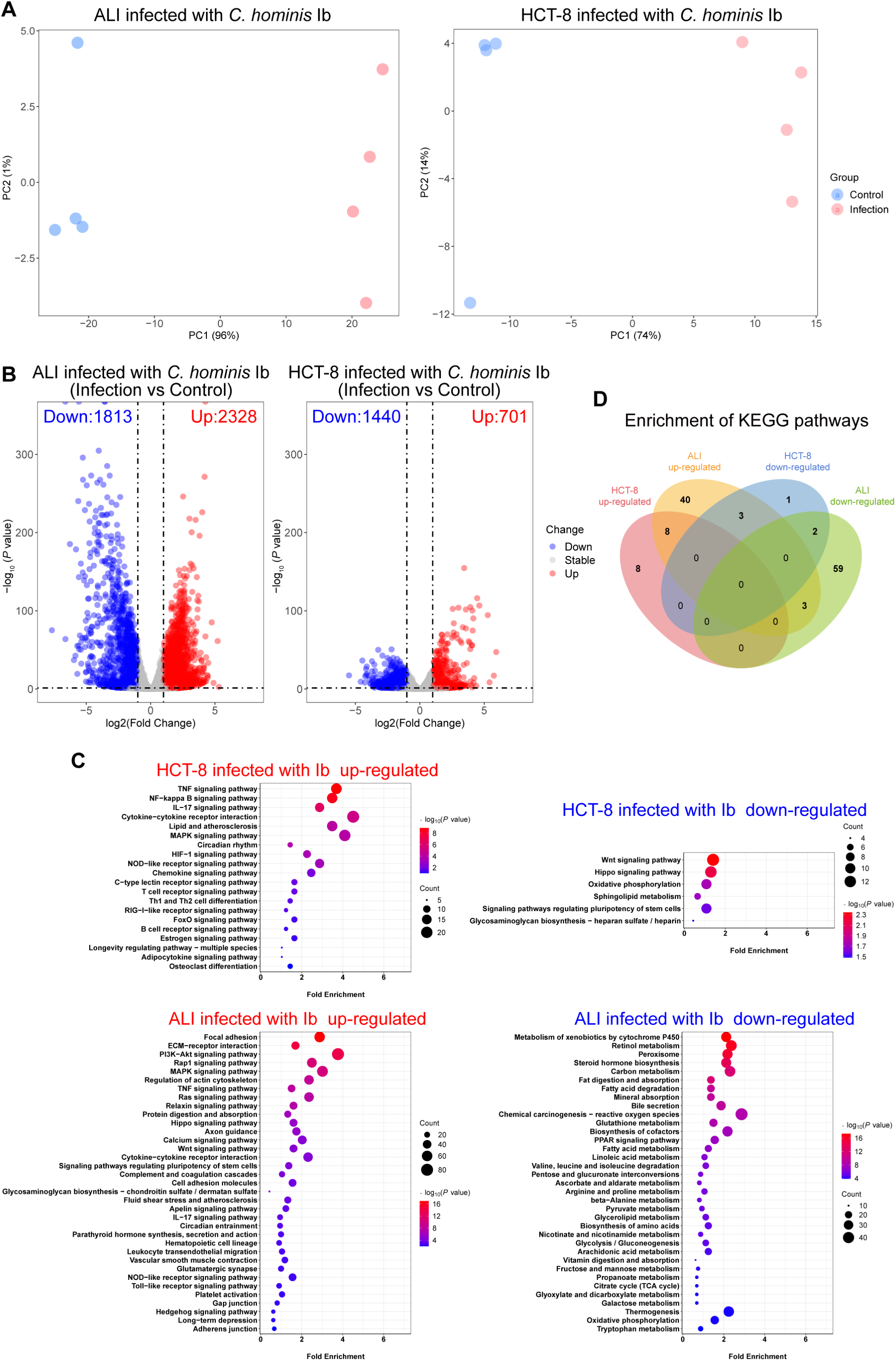
Transcriptomic responses of host genes in air-liquid interface (ALI) monolayers and HCT-8 cells after infection with *Cryptosporidium hominis*. (A) Differences in transcriptomic responses between control ALI or HCT-8 cultures and those infected with the *C. hominis* Ib subtype for 3 days, as indicated by principal component analysis of the RNA-seq data, n = 4. (B) Volcano plots showing numbers of differentially expressed genes (DEGs) between control and infected ALI or HCT-8 cultures. (C) Kyoto Encyclopedia of Genes and Genomes (KEGG) analysis of DEGs of ALI or HCT-8 cultures after infection with Ib. The y-axis shows significantly enriched KEGG pathways while the x-axis indicates the fold enrichment of DEGs. (D) Venn diagram of shared and unique KEGG pathways between ALI and HCT-8 cultures infected with Ib.

The up-regulated transcripts in the HCT-8 cells infected with Ib were significantly enriched for immune responses such as TNF, NF-κB, IL-17 and MAPK signaling pathways and NOD-like, TOLL-like and RIG-I-like receptor signaling pathways. The down-regulated DEGs involved seven pathways, including the Wnt and Hippo signaling pathways, metabolic pathways, oxidative phosphorylation, sphingolipid metabolism, pluripotency of stem cells, and glycosaminoglycan biosynthesis (Fig. 6C). In GO analysis of the RNA-seq data, the up-regulated genes were mainly involved in inflammatory and cell development processes, while the down-regulated DEGs were mainly involved in cell cycle (Fig. S2).

Compared with those in HCT-8 cells, transcriptional responses in ALI cultures infected with Ib were much more robust (Fig. 6D). Among the up-regulated DEGs in ALI cultures, most of them involved immune (including PI3K-Akt, Wnt, Ras, MAPK, TNF, and IL-17 signaling pathways and NOD-like and TOLL-like receptor signaling pathways) and inflammatory (including complement and coagulation cascades, leukocyte migration, vascular contraction and platelet activation) responses and cell adhesion and junction (including focal adhesion, ECM-receptor interaction, axon guidance, cell adhesion molecules, gap junction and adherens junction) (Fig. 6C). In contrast, the down-regulated DEGs were mainly involved in absorption, biosynthesis and metabolism of various nutrients (Fig. 6C). This was also supported by GO analysis of the DEGs (Fig. S2).

Among the 99 KEGG pathways uniquely enriched in ALI cultures, 40 pathways were up-regulated and 59 down-regulated (Table S1). In contrast, only 9 pathways were uniquely enriched in HCT-8 cells infected with Ib (including 8 up- and 1 down-regulated pathways) (Table S1). Most of the uniquely up-regulated pathways in Ib-infected ALI cultures and HCT-8 cells involved immune and inflammatory responses, while the down-regulated pathways mainly involved nutrient biosynthesis. Cell adhesion and junction and metabolism were uniquely enriched among the up- or down-regulated DEGs in ALI cultures, indicating that these cell responses were associated with the lack of further development of the parasites in HCT-8 cells. Of particular interest is the expression of the Wnt and Hippo signaling pathways, which were down-regulated in HCT-8 cultures but up-regulated in ALI cultures after *C. hominis* infection (Table S1).

### Comparative transcriptomic analysis revealed major differences in the expression of *C. hominis* genes between ALI and HCT-8 cultures

In addition, we compared the transcriptomic responses of *C. hominis* genes in HCT-8 and ALI cultures (Fig. 7A and Table S2). Among the 3913 protein-encoding genes in the *C. hominis* genomes, 1827 were expressed in both ALI and HCT-8 cultures. An additional 2005 genes had expression in ALI cultures (Fig. 7B and Table S2). Among the top 200 highly expressed genes in ALI cultures, 55 had low expression in HCT-8 cultures (log_2_FPKM <4.0). Most of them were genes encoding ribosomal proteins and *Cryptosporidium*-specific hypothetical proteins, which are required for parasites growth and proliferation. This suggests the occurrence of developmental arrest of *C. hominis* in HCT-8 cultures at 72 h (Fig. 7B and Table S2). In contrast, the KEGG analysis of the 2005 *C. hominis* genes uniquely expressed in ALI cultures had shown significant enrichment for metabolism and biosynthesis, homologous recombination and meiosis, proteolysis, and RNA degradation (Fig. S3). In particular, the expression of genes encoding oocyst wall and crystalloid body proteins was higher in Ib-infected ALI cultures than Ib-infected HCT-8 cultures (Fig. 7C). As these genes are preferentially expressed in the macrogamonts, zygotes and oocysts (12, 21, 22), these data support the generation of oocysts in ALI cultures but not in HCT-8 cultures.

**FIG 7.**
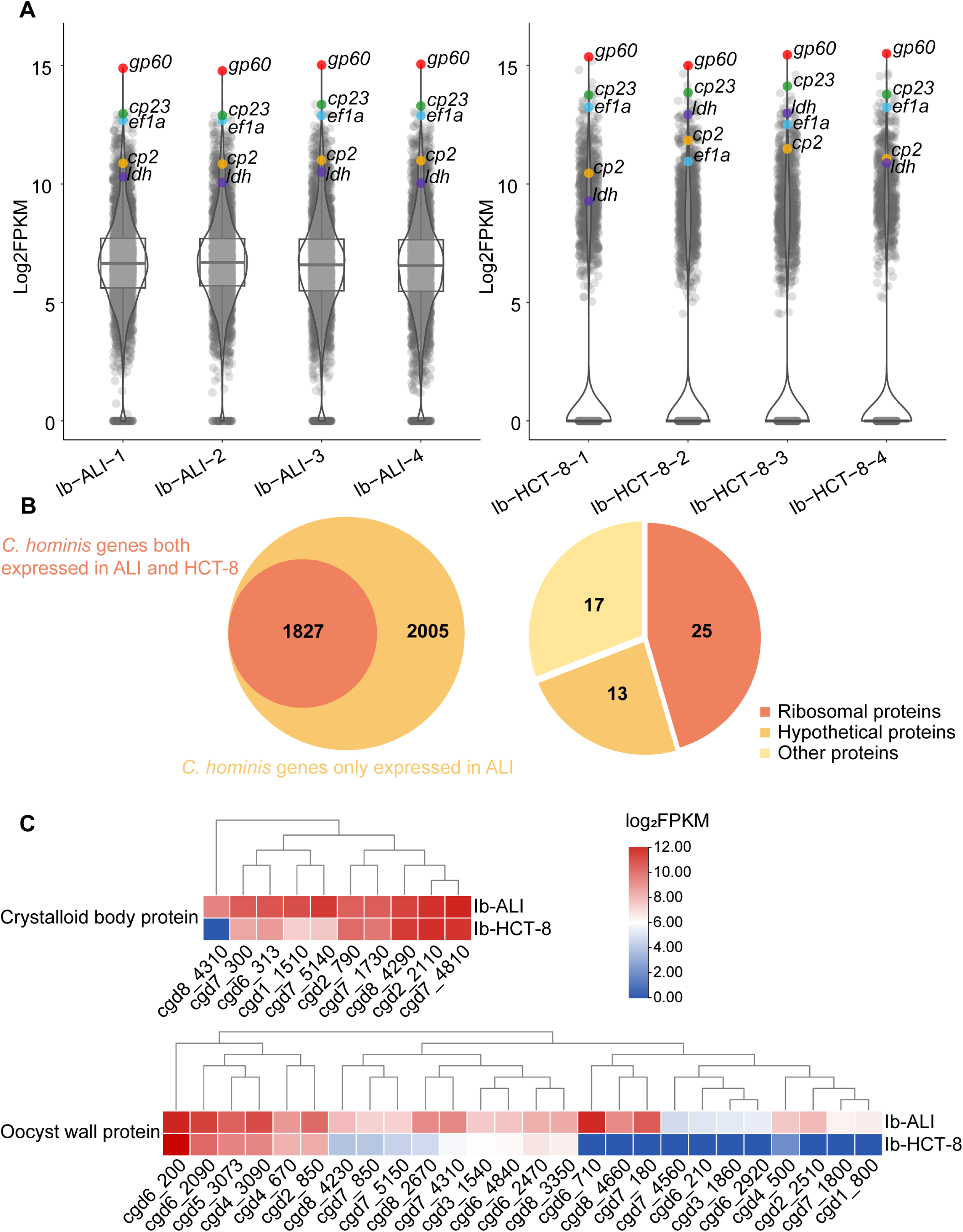
Profiles of the expression of *Cryptosporidium hominis* genes in the enteroid-based air-liquid interface (ALI) and HCT-8 cultures. (A) Violin plots of the expression profiles of *C. hominis* genes in Ib-infected ALI or HCT-8 cultures for 3 days, n = 4 for each group. (B) Venn diagram (the left figure) showing the shared (orange part) and unique genes (yellow part) of *C. hominis* Ib subtype in ALI cultures in comparison with in HCT-8 cells; pie chart (the right figure) showing the category of the 55 most differentially expressed genes (DEGs) of *C. hominis* in ALI cultures and in HCT-8 cells. The 55 genes had low expression in HCT-8 cultures (log_2_FPKM < 4.0), but were among the top 200 highly expressed *C. hominis* genes in ALI cultures. (C) Heat maps of the expression profiles of the genes encoding oocyst wall proteins (preferentially expressed in macrogamonts, zygotes and oocysts) and crystalloid body proteins (preferentially expressed in sporozoites) of *C. hominis* Ib subtype in ALI cultures and HCT-8 cells.

## DISCUSSION

In this report, we have described procedures for the generation of murine enteroids and refined the procedures for the maintenance of the ALI culture system. The modified ALI system allows the completion of the entire life cycle of *C. parvum* and *C. hominis* and propagation of isolates, producing infectious oocysts within three days. As the ALI system could be infected with frozen *C. parvum* cultures, it provides an accessible cryopreservation of *Cryptosporidium* spp. for long-term storage. RNA-seq analyses of the ALI and HCT-8 cultures infected with *C. hominis* showed significant differences in the transcriptome profiles of both the host cells and parasites, providing important information on host-parasite interactions.

The procedures of murine enteroid generation and differentiation described here allow routine cultivation of *Cryptosporidium* spp. While 3D culture systems have allowed the completion of the full life cycle of *C. parvum* in HCT-8 cells (23, 24), the newly established ALI system of *C. parvum* has the advantage of mimicking the native intestinal environment by having diverse populations of intestinal cells in cultures (17). In the present study, we have refined the operational procedures for isolation of intestinal crypts and cultivation of enteroids by modifying the treatment of i3T3 cells, culture medium, and digestion method of enteroids. In particular, the replacement of irradiation of i3T3 cells with mitomycin C treatment makes the generation and cultivation of enteroids more adaptable to most laboratories.

The modified ALI system can be adapted for in vitro studies of diverse *C. parvum* isolates. In the present study, we have used it effectively in the cultivation of one *C. parvum* IId isolate. Prior to this, in vitro studies of *C. parvum* were done exclusively with IIa subtypes (16, 17, 23, 24). Although IIa and IId subtypes differ from each other in distribution and host ranges (7), we have poor understanding of the determinants of these biological differences. In the present study, we have used the ALI system effectively in the cultivation of the virulent IIdA20G1-HLJ isolate. The oocysts generated by the enteroids were infective to mice. In fact, the IId isolate generated much higher parasite loads in ALI cultures than the reference IIaA17G2R1-Waterborne isolate. This is in agreement with the result of comparisons of the infectivity and virulence of the two isolates in GKO mice (25). Therefore, the IId-HLJ isolate was used in subsequent studies, including those on the cryopreservation of *C. parvum*. However, there could be intrinsic differences in culture characteristics among *C. parvum* isolates. IIdA20G1-HLJ oocysts produced in ALI cultures appear to be unable to excyst spontaneously, therefore failed in continuously infecting enteroids. This is different from the findings by Wilke *et al* (17).

More importantly, we have demonstrated for the first time that *C. hominis* can be cultivated in vitro using the ALI system. *C. hominis* is a primate-adapted species and a major cause of human cryptosporidiosis worldwide (4). Because of the host adaptation and difficulties in sample acquisition, no in vitro studies of *C. hominis* have been conducted thus far. The three *C. hominis* isolates we used were from crab-eating macaques in China, but have more subtelomeric genes than the common *C. hominis* subtype families (Ia, Ib, Id, Ie, and If) in humans. Because these subtelomeric genes have been associated with broad host range of *C. parvum* (26, 27), the three isolates could have broader host range than regular *C. hominis* isolates. This is probably the reason for the high infectivity of the three *C. hominis* isolates to GKO mice. Therefore, taking advantage of the infectivity of the three *C. hominis* subtypes to GKO mice, we have been able to adopt the ALI system for the cultivation of *C. hominis*. Here we have demonstrated the complete development of three *C. hominis* isolates in vitro, producing oocysts that are infective to GKO mice. Among the three *C. hominis* subtype families we used in this study, Ib and In generated higher infectivity and oocyst output than Im, re-affirming the utility of the ALI system in comparative studies of diverse *Cryptosporidium* isolates in vitro.

The ALI-*C. hominis* infection model appears to be a good mechanism for studying the host-parasite interactions in vitro. Results of the RNA-seq analysis of ALI cultures indicate that *C. hominis* infection upregulates various signaling pathways associated with immune and inflammatory responses. The intensity and nature of these host cell responses differ significantly from those induced by *C. hominis* infection of HCT-8 cells. This is expected, as the latter consists of one single type of adenocarcinoma cells of the colon compared with diverse populations of intestinal cells of the enteroids. The host cell responses detected in *C. hominis*-infected ALI cultures, however, are similar to those seen recently in *C. parvum* and *C. tyzzeri*-infected ileum (28). The RNA-seq analysis has further shown significant differences in the expression of *C. hominis* genes between ALI and HCT-8 cultures, with the former having a significantly higher expression of genes encoding proteins associated with oocysts (such as oocyst wall proteins) and sporozoites (such as crystalloid body proteins) (21). Although many genes encoding oocyst wall proteins and crystalloid body proteins have been identified in late developmental stages, we still have a rudimentary understanding of the sexual stages of *Cryptosporidium* spp. Parasite DEGs between ALI cultures and HCT-8 cells are potential candidate genes involved in the fertilization process of *Cryptosporidium* spp. (Fig. 7C). These data provide further evidence for complete development of *C. parvum* and *C. hominis* in ALI cultures.

Interestingly, the transcriptomic profiling of host cell responses has shown significant downregulation of various metabolic pathways in *C. hominis*-infected ALI cultures, including both nutrient digestion, absorption and biosynthesis. Although cryptosporidiosis is well known for causing malnutrition in children, the mechanism for this is not clear (29). The data generated here indicate that *Cryptosporidium* spp. could cause malnutrition by inhibiting host metabolism in addition to affecting nutrient absorption. This agrees with the observation in the present study of reduced number and elongation of microvilli on the epithelial cells infected with *Cryptosporidium* spp., which was previously observed in *C. parvum-*infected HCT-8 cells (30). Transcriptomic studies of *C. parvum*-infected HCT-8 cultures, however, have failed to show downregulated nutrient absorption and metabolism in host cells (31), reaffirming the conclusion that the ALI system is much better than the HCT-8 system for studying cryptosporidiosis-associated host cell responses.

The new ALI system can also be used in cryopreservation of *Cryptosporidium* spp. Reliable cryopreservation of *Cryptosporidium* oocysts has been a long-standing challenge. Although cryopreservation methods of *C. parvum* and *C. hominis* by freezing permeabilized oocysts in silica microcapillaries or ‘vitrification cassettes’ have been developed (18-20), they have not been widely used due to the limited volume of microcapillaries and inaccessibility of vitrification cassettes. In addition, frozen storage of intracellular stages of *Cryptosporidium* spp. has been tried previously (32-35). Paziewska-Harris *et al.* demonstrated that *C. parvum* could be recovered from infected cells after 6 and 12 months of storage in liquid nitrogen, with the number of recovered parasites increasing in HCT-8 cultures (35). However, the recovered parasites are unlikely to develop into oocysts, because it is well established that HCT-8 and other cell lines only support the development of *C. parvum* into sexual stages (12). Taking advantage of the ability by *C. parvum* to complete full life cycle and producing infectious oocysts in the enteroid-based ALI cultures, we have developed a novel cryopreservation method by freezing young intracellular parasites cultured in ALI or HCT-8 cultures and recovering them through further cultivation in ALI monolayers. This allows the recovered parasites to complete their development after the cryopreservation. To reduce the cost of time and labor, we amended the protocol by using HCT-8 cultures to support the growth of parasites to trophozoites before freezing. This is expected to make no difference because *C. parvum* develops equally well in both HCT-8 cells or ALI cultures during the early cultivation period.

We think the cryopreservation of trophozoites soon after the invasion of *C. parvum* is key to the success of this new approach, allowing further replication of the parasites in ALI cultures after inoculation of the cryopreserved materials. We have tried to freeze cells infected with parasites for 24 h or 48 h in preliminary experiments, but did not observe any oocysts in the ALI cultures inoculated with the recovered cells using IFA. In contrast, as shown in the study, the cryopreserved cell cultures containing early stages (ALI/HCT-6h and ALI-12h) both induced the generation of oocyst after their revival in ALI cultures. We also observed that the growth of recovered parasites of ALI-6h was higher than that of ALI-12h in ALI cultures (Fig. 4B). Because *C. parvum* in ALI-6h cultures were mostly trophozoites, we speculate that the revived trophozoites have the advantage of more rounds of asexual replication than merozoites and other stages in later cultures. Therefore, we used ALI-6h instead of ALI-12h for mouse infection.

In conclusion, the newly established enteroid system has been adapted effectively for the cultivation of host-adapted *C. parvum* and *C. hominis*, producing oocysts that are infective to mice. This allows not only comparative studies of the biology of the two major human pathogens but also cryopreservation of isolates. The use of the system in transcriptomic studies of *C. hominis* has revealed major insight into host-pathogen interactions, especially roles of the downregulated host metabolism in the induction of malnutrition by cryptosporidiosis, and candidate genes potentially involved in the regulation of sexual reproduction and oocyst development.

## MATERIALS AND METHODS

### Ethics statement

The study was conducted in compliance with the Guide for the Care and Use of Laboratory Animals. The research protocol was approved by the Committee on the Ethic Use of Animals in Research, South China Agricultural University.

### Cryptosporidium isolates

The IIa-Waterborne isolate (designated as the IOWA isolate but belonged to the IIaA17G2R1 subtype) of *C. parvum* was purchased from Waterborne, Inc. The IId-HLJ isolate of the IIdA20G1 subtype was initially obtained from a dairy calf on a farm in Heilongjiang (HLJ), China and maintained through passages in GKO mice. This isolate has higher infectivity and some unique genomic sequence characteristics in comparison with the IIaA17G2R1-Waterborne isolate as previously described (25). Oocysts of nonhuman primate-adapted *C. hominis* isolates of the IbA12G3, ImA20, InA17 subtypes were initially obtained from crab-eating macaques (*Macaca fascicularis*) in China using sucrose and cesium chloride gradient centrifugations (36, 37). In a separate study, comparative genomic analysis of the three macaque *C. hominis* isolates and common *C. hominis* isolates from humans confirmed their *C. hominis* identity, with 16315-16709 SNPs from common *C. hominis* (compared to ∼20000 SNPs between the anthroponotic IIc and zoonotic IIa or IId *C. parvum*) (38). They have two subtelomeric MEDLE genes and one subtelomeric INS gene that are usually found only in *C. parvum*. In experimental infection of GKO mice, the three macaque isolates induced high levels of oocyst shedding (> 10^6^ OPG) and long oocyst shedding duration (> 80 days) (38). These three unique *C. hominis* isolates have been maintained in our laboratory through passages in GKO mice since their initial isolation in August 2020, with subtype confirmation after each passage. All *C. parvum* and *C. hominis* oocysts obtained were stored at 4°C in PBS containing antibiotics for less than two months before being used.

### Establishment and maintenance of enteroids

Ileum crypts were isolated from 6-week-old female C57BL/6 mice. Two to three sections (∼4 cm) were harvested from the small intestine of sacrificed mice, opened longitudinally, and washed with ice-cold PBS three times. They were cut into 2∼3 mm pieces and rotated in a 15 ml tube at 4°C and 20 rpm in EDTA chelation buffer (ice-cold PBS containing 2 mM EDTA and 100 mg/ml gentamicin) for 40 min. The tissue fragments were allowed to settle on ice for 30 sec. After replacing the buffer with ice-cold PBS (∼5 ml), the tube was shaken violently until the supernatant was clouded. The tissue suspension was filtered through a pre-wetted 70 μm filter (Corning) to collect the filtrate. This procedure was repeated 3∼4 times, and the pooled filtrate was centrifuged at 4°C and 300 g for 5 min. Crypts in the tube were collected after removing the supernatant, and resuspended in 50% L-WRN conditioned medium (CM, Table S3) mixed 1:1 with Matrigel (BD Biosciences). The crypt suspension (55 μl per well) was transferred into a pre-warmed 24-well plate, and incubated at 37°C for 10 min to congeal the Matrigel. Afterwards, 700 μl CM was added into each well gently, not disturbing the Matrigel domes. The plate was incubated at 37°C and 5% CO_2_, with the medium being replaced with fresh CM every 2∼3 days. The enteroids were passaged after 7 days by the addition of 1 ml Gentle Cell Dissociation Reagent (STEMCELL) and repeated pipetting to break up the enteroid-Matrigel domes. The suspension was digested at room temperature for 10 min and centrifuged at 4°C and 300 g for 5 min. After removing the supernatant, the enteroids were washed with 1 ml DMEM/F-12 by centrifugation. Aliquots of the enteroids were mixed with Matrigel, plated into a pre-warmed 24-well plate, and cultured as described above. We passaged enteroids using the 1:4 split ratio every 3 or 4 days.

NIH/3T3 embryonic mouse fibroblast cells (ATCC, CRL-1658) were cultured at 37°C and 5% CO_2_ in DMEM (Gibco) supplemented with 10% fetal bovine serum (FBS, Gibco) and 1U/ml penicillin/streptomycin, and passaged every 2 days in a 1:5 split.

### Generation of ALI culture system and infection with Cryptosporidium spp

The procedures described in the previous study (17) were used in the generation of the ALI culture system, with the following modifications: 1) we treated the i3T3 cells with mitomycin C instead of irradiation; 2) we changed the digestion of enteroids from the use of trypsin solution to a more gentle approach (using TrypLE Express) to reduce damages to the enteroid cells; 3) we supplemented the CM with additional growth factors and antibiotics such as mouse EGF and Primocin (Table S3).

For the construction of the ALI system, 3T3 cells were incubated in 4 μg/ml mitomycin C (CST, cat# 51854S) for 1 h and washed three times with PBS to create i3T3 cells as the feeder cells. The i3T3 cells were plated at the density of 8×10^4^ i3T3 cells on a Transwell insert (polyestermembrane with 0.4 μm pores; Biofil, 12 Transwells per 24-well plate) that was pre-coated with diluted Matrigel (Matrigel: ice-cold PBS=1:10). DMEM with 10% FBS and 1U/ml penicillin/streptomycin was added to the top (100 μl) and bottom (400 μl) parts of the Transwell and incubated at 37°C and 5% CO_2_ for approximately 24 h. The next day, the enteroids were trypsinized using TrypLE Express (Gibco) and roughly pipetted. After 10 min digestion at room temperature, the cells were centrifuged at 300 g for 5 min and washed with DMEM/F-12. The cells were resuspended in 1 ml CM and filtered through a pre-wetted 40 μm filter (Corning) to remove undigested enteroids, and seeded on the i3T3 cell layers with 5×10^4^ cells per Transwell. The medium in the top (200 μl) and bottom (400 μl) parts of Transwell was changed every 2 days as described above. To create the ALI for cell differentiation, the medium in the top chamber of the Transwell was removed at day 7 of the culture, while the medium in the bottom chamber was changed every 2 days. Three days after removing the top medium, the ALI system was infected with oocysts or purified sporozoites as described previously (17) (Fig. 1A). Sporozoites were purified from excystation suspension of oocysts using 3 µm filters (Whatman).

### Cryopreservation of C. parvum-infected ALI or HCT-8 cultures

Cryopreservation was performed on ALI cultures (at 6 h or 12 h) or HCT-8 cells (at 6 h) infected with sporozoites purified by using 3 µm filters. For cryopreservation, cells from 2∼3 Transwells (for ALI cultures) or one well of 48-well plates (for HCT-8 cultures) were placed into each cryovial (1 ml). Prior to the collection of culture materials, the culture medium was replaced with ice-cold PBS, and the cultures infected with *C. parvum* were pipetted repeatedly. After centrifugation at 4°C and 300 g for 5 min and one wash with DMEM/F-12, the cells harvested were resuspended in cryovials with 1 ml of cryoprotectant (STEMCELL) for the cryopreservation of organoids. After storage in a freezing container at -80°C for 24 h, the cryovials were transferred to liquid nitrogen for long-term storage.

For thawing the frozen cultures, cryovials were placed in a 37°C water bath for 2∼2.5 min. When the freezing medium became liquid, the cells infected with parasites were transferred immediately to 15-ml tubes containing 3 ml DMEM/F-12 with 1% BSA (Biofroxx). After careful mixing, the cell suspension was centrifuged at 4°C and 200 g for 5 min to remove the cryoprotectant. After removing the supernatant, the intracellular parasites were resuspended with 0.5 ml CM and seeded into 1∼2 Transwells in ALI cultures. The infected ALI cultures were maintained as described above.

### Measurement of Cryptosporidium growth

In measurement of *Cryptosporidium* growth in the ALI system, three wells of cultures were used for each infection group at each sampling point. DNA was extracted from each well using the QIAamp DNA Mini kit (Qiagen) after five freeze-thaw cycles in liquid-nitrogen and 56°C water-bath. In measurement of *Cryptosporidium* shedding in GKO mice, DNA was extracted from 0.1 g fecal pellets from each mouse using the FastDNA SPIN Kit for Soil (MP Biomedicals). The infection intensity in Transwells and mice was assessed using SYBR Green-based 18S-LC2 qPCR as described (39). Student’s *t*-test was used to compare the parasite burden between groups. Differences were considered significant at *P* < 0.05.

### Microscopic analyses of ALI cultures

For immunostaining, monolayers in the Transwell inserts were fixed with 0.2 ml 4% paraformaldehyde for 10 min, permeabilized with 0.1% Triton X-100 (Sigma) for 10 min, and blocked with 1% BSA for 60 min. The cells were then incubated at room temperature with Crypt-a-Glo (Waterborne, 1:20) and Sporo-Glo (Waterborne, 1:20) for 60 min. DAPI (CST, DAPI: PBS = 1:2 000) was used to stain the nuclear DNA. The membrane of the Transwell insert was cut with a scalpel and plated on a glass slide for microscopy.

For histological analyses, Transwells were fixed at room temperature by adding 4% paraformaldehyde into both the top (0.2 ml) and bottom (0.4 ml) chambers for 30 min. The Transwell membranes were cut from the inserts using a scalpel and processed for paraffin embedding. Sections of 4 μm in thickness were cut and stained with conventional procedures for the hematoxylin and eosin (H&E) staining and immunohistochemistry. The primary antibodies used in immunofluorescence analysis included mouse anti-villin (1:1000, Abcam), rabbit anti-mucin 2 (1:1000, Abcam) and rabbit anti-Ki-67 (1:500, Servicebio), while the secondary antibodies included goat anti-mouse IgG Alexa Fluor 488 (Abcam) and goat anti-rabbit IgG Alexa Fluor 568 (Invitrogen).

The slides generated were examined under an Olympus BX53 (Olympus, Japan) fluorescence microscope. Images were acquired using the CellSens Standard1 software (Olympus) and processed in ImageJ or Photoshop.

For ultrastructural analyses, ALI monolayers were fixed with 2.5% glutaraldehyde and processed for SEM and TEM using conventional procedures. They were examined under an EVO MA 15/LS 15 (Carl Zeiss Microscopy GmbH, Jena, Germany) and Talos L120C (Thermo Fisher, Waltham, USA), respectively.

### Experimental infection of GKO mice with cultured parasites

Specific-pathogen-free GKO mice with a C57BL/6 background were used in assessment of the infectivity of oocysts produced in culture. They were housed in individually and supplied with sterilized diet, water, and bedding. The experiment was conducted with GKO mice of four weeks of age.

For the assessment of the infectivity of oocysts produced from ALI system without cryopreservation, four or three GKO mice (n = 3 or 4) were orally gavaged with 200 μl of bleach-treated ALI material (equivalent to 1.25 Transwells per mouse) harvested one (D1) or three (D3) days after the infection with sporozoites. Fecal samples were collected from them every three days. For the assessment of the infectivity of oocysts produced from the ALI system after cryopreservation, GKO mice were pretreated with 1 mg/ml ampicillin (Macklin), 1 mg/ml streptomycin (Macklin) and 0.5 mg/ml vancomycin (Macklin) for three days before infection. Each mouse in the infection group (n = 3) was infected with cell materials of 2∼3 Transwells, which contained ALI cultures infected with recovered parasites for 4 days. In contrast, mice in the control group (n = 3) were gavaged with PBS. Fecal samples were collected from each mouse every three days for the measurement of oocyst shedding using a 18S-LC2 qPCR as described above. The subtype identity of the oocysts shedding in GKO mice was confirmed by sequence analysis of the *gp60* gene as described (40).

### RNA-seq analysis of C. hominis-infected ALI and HCT-8 cultures

RNA was isolated from uninfected and *C. hominis*-infected Transwell cultures or HCT-8 cells three days after the infection. To obtain sufficient RNA for sequencing, two Transwells were combined as one sample (n = 1), and four samples (n = 4) each of the infected and control cultures were used in the RNA-Seq analysis. For this, 0.5 ml ice-cold PBS was used to harvest the monolayers by centrifugation at 4°C and 300 g for 5 min. After removing the supernatant, 0.2 ml of Trizol was added into each sample and pipetted several times until the suspension became opaque. The samples were frozen in liquid-nitrogen immediately and stored at -80°C before RNA-seq using the Illumina NovaSeq 6000 at Guangzhou Genedenovo Biotechnology Co. Transcriptomic analysis of the RNA-seq data for host cell responses and *C. hominis* gene expression was performed as described previously (28). KEGG and GO analyses of DEGs were used to identify biological pathways and terms involved. The RNA-seq data generated in this study have been deposited in the SRA database of the National Center for Biotechnology Information under the accession number PRJNA988932 (https://www.ncbi.nlm.nih.gov/sra/PRJNA988932).

## SUPPLEMENTALMATERIAL

Supplemental material is available online only.

**FIG S1**, TIF file, 6.48 MB. **FIG S2,** TIF file, 2.11 MB. **FIG S3**, TIF file, 1.46 MB.

**Table S1**, **Table S2** and **Table S3**, XLSX file, 0.55 MB.

## ACKNOWLEDGEMENTS

We thank Yingxin Liang for assistance in the generation of murine enteroids, Ruilian Jia, Ni Huang, Xuehua Chen, Haoyu Chen and Huimin Liu for provision of laboratory mice, and staff at the South China Agricultural University microscopy core facility for assistance with electron microscopy.

This study was funded by National Natural Science Foundation of China (32150710530, 31972697 and 32030109), Double First-Class Discipline Promotion Project (2023B10564003), 111 Project (D20008), and Innovation Team Project of Guangdong University (2019KCXTD001).

We declare that we have no conflicts of interest.

## FIGURE LEGENDS

**FIG S1** Histological examination of air-liquid interface (ALI) cultures and mice infected with cryopreserved IId subtype of *Cryptosporidium parvum*. The images were taken from the sections after staining with H&E or rabbit antibodies against *C. parvum* (IHC). (A) Sections of new ALI cultures infected with cryopreserved ALI cultures (which were infected with *C. parvum* for 6 h or 12 h and then cryopreserved in liquid nitrogen: ALI-6h and ALI-12h) for 3 days. Scale bars, 10 μm. (B) Sections of ALI cultures infected with cryopreserved HCT-8 cultures (which were infected with *C. parvum* for 6 h and then cryopreserved in liquid nitrogen: HCT-6h) for 3 days. Scale bars, 10 μm. (C) Images of H&E-stained sections of the ileum from mice inoculated with bleach-treated ALI cultures that were infected with cryopreserved ALI-6h and HCT-6h described above. The mice were sacrificed 15 days after the infection. Scale bars, 20 μm.

**FIG S2** Gene ontology (GO) analysis of differentially expressed genes (DEGs) of the host cell in the air-liquid interface (ALI) monolayers (a) or HCT-8 cells (b) infected with *Cryptosporidium hominis*. The y-axis shows the enriched GO terms while the x-axis indicates the fold enrichment of DEGs. Both the up-regulated (left) and down-regulated (right) responses are shown.

**FIG S3** Kyoto Encyclopedia of Genes and Genomes (KEGG) enrichment analysis of 2005 *Cryptosporidium hominis* genes uniquely expressed in the air-liquid interface cultures in comparison with HCT-8 cultures. The y-axis shows the enriched KEGG pathways, while the x-axis indicates the fold enrichment.

